# Mechanical and Microstructural Properties of Pediatric Anterior Cruciate Ligaments and Autograft Tendons used for Reconstruction

**DOI:** 10.1101/338905

**Authors:** Elaine C Schmidt, Matthew Chin, Julien T Aoyama, Theodore J Ganely, Kevin G Shea, Michael W Hast

## Abstract

**Background:** Over the last several decades there has been a steady increase in pediatric ACL tears, particularly in young female basketball and soccer players. Because allograft tissue for pediatric ACL reconstruction (ACLR) has shown high rates of failure, autograft tissue may be the best option for ACLR in this population. However, the differences in structure and mechanical behavior of these tissues are not clear.

**Purpose:** This study sought to characterize mechanical and microstructural properties in pediatric ACLs and autograft tissues using a rare cadaveric cohort (mean age 9.2 years).

**Study Design:** Descriptive laboratory study.

**Methods:** ACLs, patellar tendons, quadriceps tendons, semitendinosus tendons, and iliotibial bands (ITBs) were harvested from five fresh-frozen pediatric knee specimens (3M, 2F) and subjected to a tensile loading protocol. A subset of contralateral tissues were analyzed using brightfield, polarized light, and transmission electron microscopy.

**Results:** Patellar tendons exhibited values for ultimate stress (5.2±3.1 MPa), ultimate strain (35.3±12.5%), and Young’s Modulus (27.0±8.0 MPa) that were most similar to the ACL (5.2±2.2 MPa; 31.4±9.9%; 23.6±15.5 MPa). Semitendinosus tendons and ITBs were stronger but less compliant than the quadriceps or patellar tendons. ITBs exhibited crimp wavelengths (24.3±3.1 um) and collagen fibril diameters (67.5±19.5 nm) that were most similar to the ACL (24.4±3.2 um; 69.7±20.3 nm).

**Conclusion:** The mechanical properties of the patellar tendon were almost identical to that of the ACL. The ITB exhibited increased strength and similar microstructure to the native ACL. These findings are not entirely congruent to studies examining adult tissues.

**Clinical Relevance:** Results suggest that ITB tissue may be the preferable choice as an autograft tissue in pediatric ACL reconstructions.

**Key Terms:** Pediatric, ACL reconstruction, mechanical properties, microstructural properties, patella tendon grafts, quadriceps tendon grafts, hamstring grafts

**What is Known about the Subject:** Due to the extreme rarity of pediatric cadaveric specimens, very little is known about these tissues.

**What this Study Adds to Existing Knowledge:** This suite of data can be used to further optimize the design and selection of grafts for reconstruction and may provide insight into the development of constitutive musculoskeletal models.

## INTRODUCTION

A large amount of research has been devoted to characterizing the mechanical and microstructural properties of the tendons and ligaments surrounding the adult human knee. Due to the extreme rarity of these cadaveric specimens, relatively little is known about these properties in the pediatric population. While the effects of senescent aging on the tensile properties of these structures have been established,^22,35,58^ extrapolating this data back to pre-pubescent ages is inadequate. Over the past several decades, the clinical need for this data in pediatric orthopedics has grown in parallel with the steady increase of diagnosed ligament tears in skeletally immature patients.^3,11,27,36^ This trend has been attributed to several factors, including increased participation in youth sports, sport specialization, year-round play, and an increase in the number of adolescents competing at higher levels of competition.^13^ Pediatric ACL tears account for the majority of these knee injuries, particularly in young female soccer and basketball athletes.^16,47^

In the pediatric patient, surgical treatment of ACL deficiency is complicated by the potential risk of injury to the physis. There is currently large debate and practice variation in initial management, operative timing, and operative technique for pediatric ACL reconstruction (ACLR). Adolescents approaching skeletal maturity can be managed similar to adults with a complete transphyseal reconstruction.^15^ However, if the patient’s physes are still open, physeal-sparing or partial transphyseal techniques are often preferred in order to prevent premature physeal closure and post-operative growth disturbance.^15,23,28^ In choosing the best course of treatment, the surgeon must consider both the pediatric patient’s bone growth potential and the need for graft stability and durability in the face of a return to sport and a longer remaining lifespan.^41^

Graft choices for ACLR include the hamstrings tendon, bone-patellar tendon-bone, quadriceps tendon, and iliotibial band (ITB). Autografts are generally preferred because allografts have been shown to have increased failure rates in younger and more active children due to slower incorporation and higher infection rate.^14^ Each of these graft options have been shown to be clinically successful, but also possess their own suite of risks. Unresolved issues regarding harvesting, biologic incorporation, and donor site morbidities remain controversial.^9,25,54,57^

There remains a dearth of knowledge about the mechanical and structural properties of autograft tissues in the pediatric population. It is currently unknown whether patellar tendon, ITB, or hamstring tendons possess the appropriate structural properties and mechanical durability to be well-suited to act as an ACL surrogate. Gaining a better understanding of these relationships will likely improve surgical outcomes in pediatric patients, particularly as ACLR procedure frequency continues to accelerate.^40^ While many in vitro studies have examined the material properties of ACLs and common grafts used for knee ligament reconstruction in adults ^48,49^ and in skeletally immature animal models,^7,10,59^ to our knowledge, no analogous studies have been conducted for the pediatric population. Therefore, the purpose of this study was to characterize the mechanical properties and microstructure of ACLs and the most common tendons used for pediatric ACLR.

## METHODS

Five fresh-frozen pediatric cadavers (3M, 2F, average age 9.2 years) were acquired through donation from Allosource (Centennial, Colorado). Due to the rarity of this cohort, portions of the knees were dissected, refrozen at −20°C, and sent to various research labs throughout the country. This study received tissue samples from all five donors, primarily for the purposes of mechanical testing (Figure 1). In some cases, tissues from the contralateral limb were also available for testing and microstructural analyses were performed on these specimens. Specifically, the contralateral ACLs from Donor 3 (male, age 10) and Donor 4 (male, age 9) were available for hematoxylin-eosin (H&E) histology, polarized light, and transmission electron microscopy (TEM) analysis. All of the contralateral tendons and ligaments from Donor 5 (female, age 9) were also available and used for these microstructural analyses. This allowed for comparison between the ACL and different candidate autograft tissues within a single subject.

**Figure 1:**
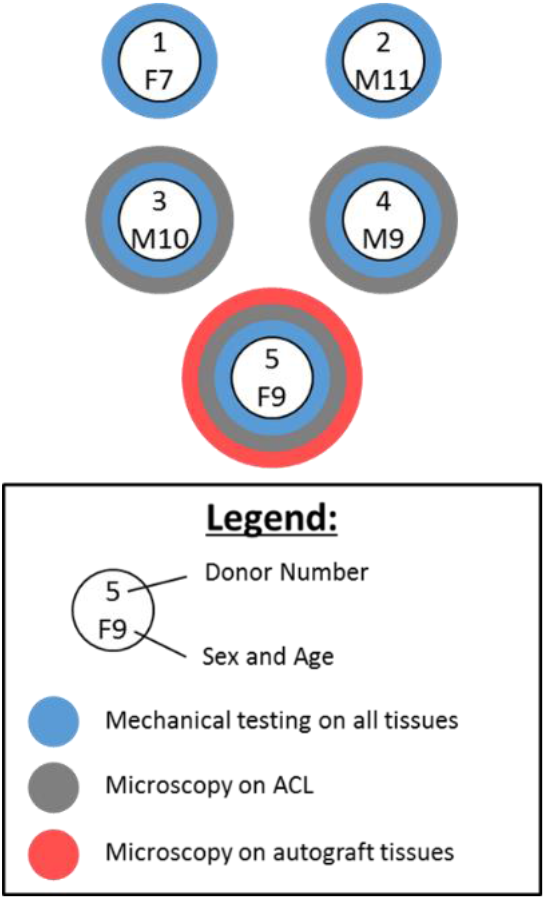
*Schematic showing testing designations for the procured pediatric knee specimens*.

### Mechanical Testing

Mechanical testing was performed on a total of 25 specimens from five donors. Specimens were prepared for testing by cutting them into standard dog bone shapes at the mid-substance (ACL, patellar tendon, quadriceps tendon; Figure 2A) or distal substance (semitendinosus and ITB) with a custom-built jig. Cross-sectional areas (CSAs) were measured along the gauge length of prepared specimens with a noncontact laser-based measurement system and averaged (Figure 2B). Specimen ends were then placed in custom aluminum clamps and attached to a 3 kN load cell on a universal testing frame (TA Instruments ElectroForce 3330, Eden Prairie, MN) to perform uniaxial tensile testing (Figure 2C&D). One ACL (Donor 1) was excluded from testing due to its small size and poor quality, which made it impossible to test. Tissues were kept hydrated throughout sample preparation and mechanical testing.

**Figure 2:**
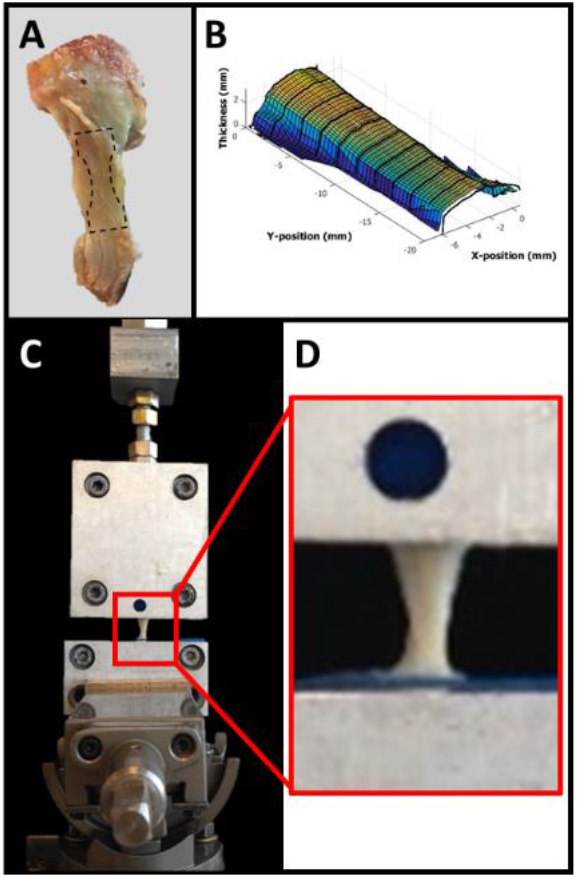
*(A) Representative image of a harvested pediatric bone-patellar tendon-bone segment. The dashed lines show the location where the specimens were cut into dog-bone shapes at the mid-substance. (B) Topographical map of the middle portion of a dog-boned specimen taken with a laser based 3D scanner. (C&D) Photograph of the tensile testing setup showing how the custom-made grips and specimens were subjected to tensile loads*.

The tensile loading protocol was based on a previously published study.^33^ Briefly, the tissue was preloaded from a slack position to 5 N and then preconditioned with 10 cycles between 10 N and 5 N at 0.4% strain rate. After dwelling at 5 N, the specimen underwent a stress-relaxation protocol followed by a ramp to failure at a constant quasistatic strain rate of 0.03% per second (Supplemental Figure 1). Failure of the specimen was confirmed mechanically by a sharp decrease in force and visually by the observance of frayed fibers at the mid substance. The viscoelastic parameter of percent relaxation was calculated as the percent change in stress from peak stress to equilibrium (Figure 2A). A bilinear constitutive model with a least squares fit ^8,29,55^ (Figure 2B; Eq. 1) was applied to the stress-strain data to quantify the moduli in the toe and linear regions, as well as the stress and strain values corresponding to the transition point:

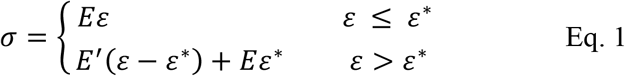

where *E* is the slope of the toe modulus, *E′* is the slope of Young’s modulus, *ε* is the strain, *ε*\* is the transition strain, and *σ* is the stress. Although the toe region is non-linear, the model produced a practical and conservative approximation of the toe modulus and transition point. Stiffness was calculated as the slope of the load-displacement curve and strain energy density, which represents the energy absorbed before failure, was calculated as the area under the stress-strain curve.

### Histology and Polarized Light Microscopy

Samples for histology were fixed in 10% neutral buffered formalin and embedded in paraffin. Serial sections (~8 μm) were stained with H&E and viewed with brightfield microscopy (Figure 3A). Ten randomly chosen regions (0.35 × 0.26 mm^2^ each) were evaluated for each specimen and mean cell density was measured using Image-J software (NIH, Bethesda, MD). Collagen crimp properties were examined using polarized light microscopy, as previously described in the literature.^30^ Crimp wavelength was determined by crossing the polarizer and analyzer at 90°, rotating the specimens until maximal extinction in the dark crimp bands occurred, and calculating the distance spanned by a dark and light band unit (Figure 3B).^12,60^ Scaled images from ten randomly chosen regions (0.35 × 0.26 mm^2^ each) were obtained and consecutive crimp wavelengths were individually measured and averaged. A minimum of 100 crimps were analyzed over the ten images for each specimen. The quality of the histology sections for one ACL (Donor 4) were determined to be too poor to be included in subsequent anaylsis and were consequently excluded.

**Figure 3:**
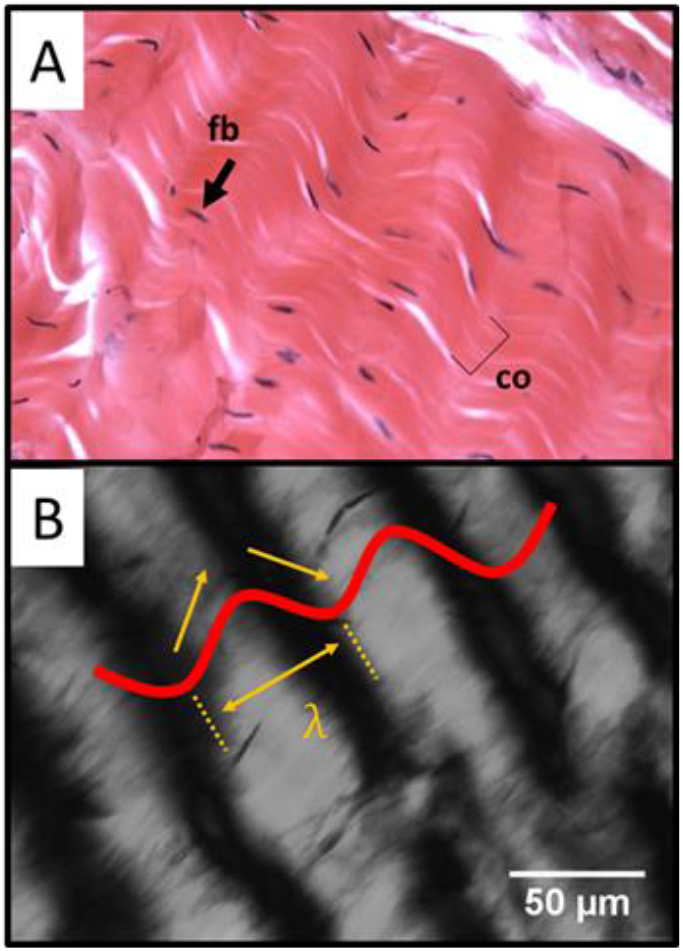
*(A) H&E stained pediatric patellar tendon specimen viewed using brightfield microscopy. Fibroblast nuclei (fb) and collagen (co) are indicated. B) Under polarized light microscopy, discrete crimps are identified and the wavelength (λ) is measured in micrometers. 20X magnification, scale bar: 50 μm*.

### Transmission Electron Microscopy

Tissues for electron microscopic examination were fixed with 2.5% glutaraldehyde, 2.0% paraformaldehyde in 0.1M sodium cacodylate buffer overnight at 4°C. After subsequent buffer washes, the samples were post-fixed in 2.0% osmium tetroxide for 1 hour at room temperature, and rinsed in DH2O prior to *en bloc* staining with 2% uranyl acetate. After dehydration through a graded ethanol series, the tissue was infiltrated and embedded in EMbed-812 (Electron Microscopy Sciences, Fort Washington, PA). Thin sections were stained with uranyl acetate and lead citrate and examined with an electron microscope (JEOL 1010, JEOL USA, Inc, Peabody, MA) fitted with a digital camera and AMT Advantage image capture software (Advanced Microscopy Techniques, Woburn, MA).

Ten micrographs were obtained at 60,000x magnification transverse to the load bearing axis for each specimen, and each micrograph was analyzed using a semi-automated protocol in Image-J/Fiji software ^44^ (Figure 4A). A machine learning classifier was trained to distinguish the darker fibrils from the lighter background of the micrograph (Figure 4B). The fibrils were then completely differentiated from the background using a binary thresholding algorithm and separated from other fibril edges using a watershed segmentation algorithm (Figure 4C). Each discrete fibril was fit with an ellipse and the Feret’s minor diameter was determined, which reduced error introduced by unintended oblique sectioning of the fibrils (Figure 4D). The distributions of fibril diameters were fit using a kernel density estimation to show the modality of their profiles. A minimum of 3,000 fibrils were analyzed over the 10 TEM micrographs for each specimen.

**Figure 4:**
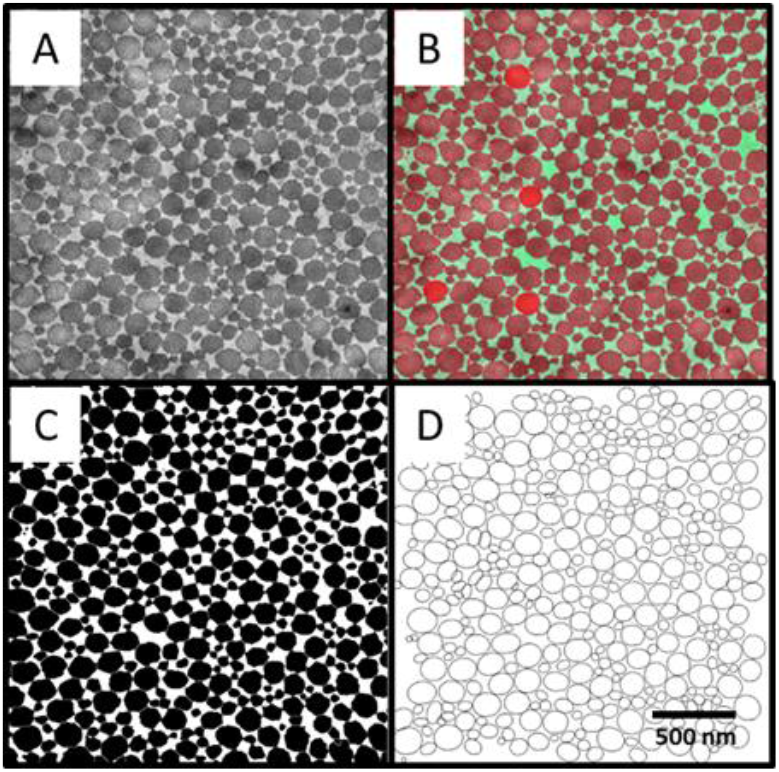
*TEM micrograph analysis protocol. (A) Micrograph for a pediatric patellar tendon specimen. (B) Machine learning segmentation of fibrils (red) from background (green). (C) Complete segregation of the fibrils using thresholding and watershed techniques. (D) Ellipse fit to each individual fiber*.

## RESULTS

### Mechanical Testing

The patellar tendon exhibited mechanical properties that were most similar to that of the ACL, particularly for ultimate stress, Young’s modulus, and strain energy density (Table 1). The toe moduli of the patellar tendon and ACL were also similar, however, the transition stresses and transition strains of the patellar and quadriceps tendons were more similar to each other than to the ACL (Table 2). Pediatric semitendinosus tendon was stronger and less compliant than the ACL and other graft candidates in addition to exhibiting considerably larger values for Young’s modulus, stiffness, and strain energy density. The patellar tendon exhibited the greatest percentage of stress-relaxation while the values for ACL, quadriceps tendon, semitendinosus tendon, and ITB were relatively similar.

**Table 1.** Results (mean ± standard deviation) for pediatric ACL and graft candidate tendons obtained from tensile testing.

|  | n | % Stress Relaxation | Ultimate Stress (MPa) | Ultimate Strain (%) | Young's Modulus (MPa) | Stiffness (N/mm) | Strain Energy Density (MPa) |
| --- | --- | --- | --- | --- | --- | --- | --- |
| ACL | 4 | 26.5 ± 9.2 | 5.2 ± 2.2 | 31.4 ± 9.9 | 23.6 ± 15.5 | 39.6 ± 17.6 | 0.8 ± 0.4 |
| Patellar | 5 | 40.6 ± 4.8 | 5.2 ± 3.1 | 35.3 ± 12.5 | 27.0 ± 8.8 | 17.5 ± 5.9 | 1.2 ± 0.9 |
| Quadriceps | 5 | 33.2 ± 9.8 | 12.1 ± 8.3 | 38.9 ± 15.7 | 61.9 ± 65.0 | 39.1 ± 31.6 | 2.5 ± 1.9 |
| Semitendinosus | 5 | 26.7 ± 4.7 | 29.0 ± 11.6 | 20.8 ± 7.2 | 197.2 ± 34.4 | 100.8 ± 17.6 | 3.1 ± 1.9 |
| ITB | 5 | 22.6 ± 3.0 | 29.0 ± 10.6 | 28.1 ± 10.6 | 161.7 ± 100.2 | 61.6 ± 22.5 | 3.8 ± 1.3 |

**Table 2.** Toe region results (mean ± standard deviation) for pediatric ACL and graft candidate tendons obtained from tensile testing.

|  | n | Transition Stress (MPa) | Transition Strain (%) | Toe Modulus (MPa) |
| --- | --- | --- | --- | --- |
| ACL | 4 | 0.8 ± 0.3 | 5.5 ± 1.9 | 13.1 ± 7.2 |
| Patella | 5 | 0.4 ± 0.2 | 2.4 ± 0.7 | 16.9 ± 6.0 |
| Quadriceps | 5 | 1.0 ± 1.1 | 2.8 ± 0.3 | 33.6 ± 37.7 |
| Semitendinosus | 5 | 3.0 ± 1.6 | 5.1 ± 2.7 | 65.2 ± 13.5 |
| ITB | 5 | 2.8 ± 0.9 | 6.2 ± 5.2 | 58.9 ± 29.5 |

### Histology, Polarized Light, and TEM

Cell density analysis of H&E stained sections showed higher concentrations of fibroblasts in the ACL and ITB samples from Donor 5 compared to the other tendons (Figure 5; Table 3). Mean fibroblast cell count in the ITB was considerably greater (1734.4 ± 327.9 cells/mm^2^) than the other candidate graft tendons, especially the quadriceps tendon (296.5 ± 49.2 cells/mm^2^).

**Figure 5:**
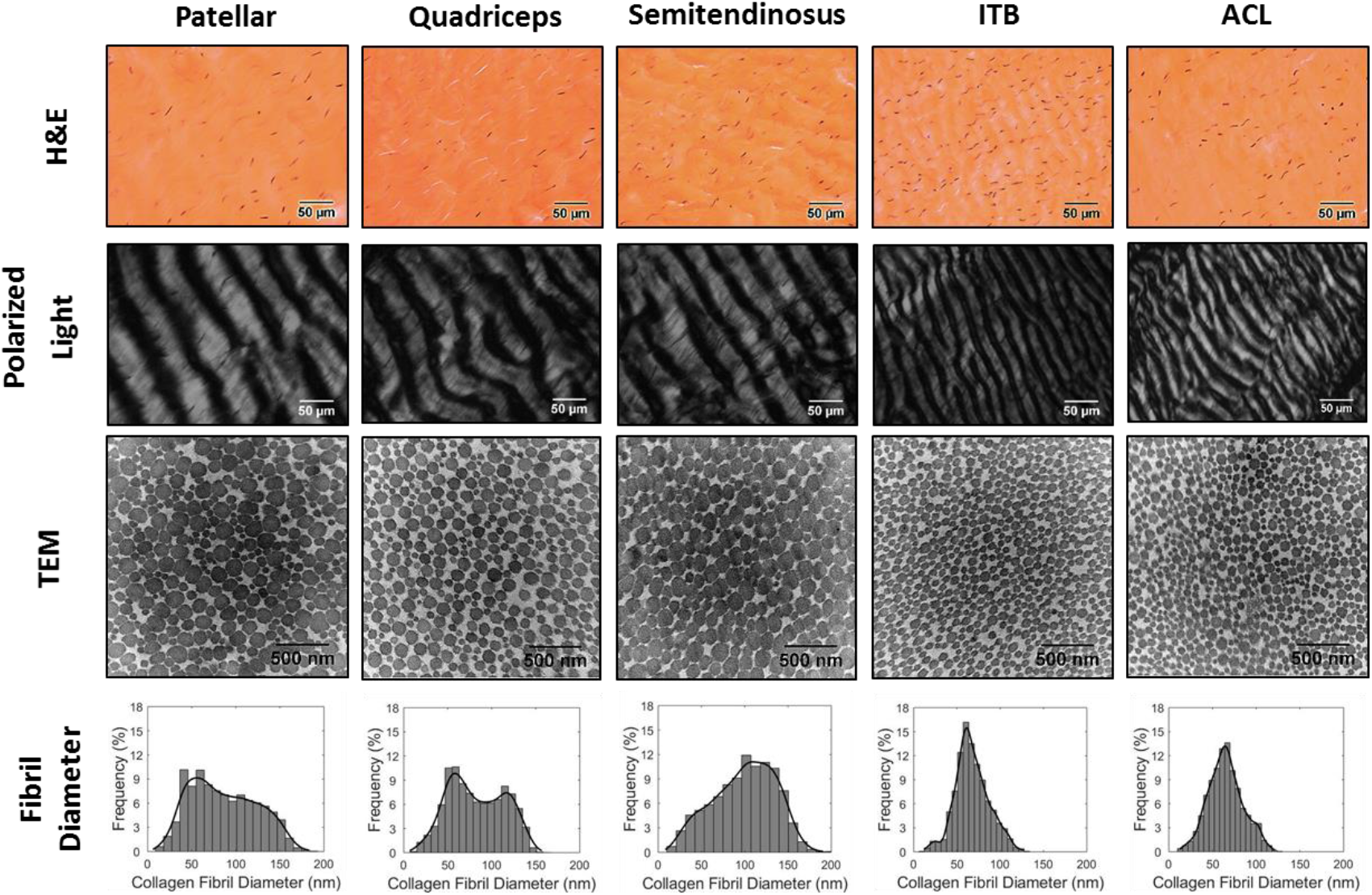
*Histology, polarized light, and TEM results for the pediatric patellar tendon, quadriceps tendon, semitendinosus tendon, ITB, and ACL samples. First row: representative H&E-stained histology sections. Second row: representative polarized light images showing crimp morphology. Third row: TEM micrographs showing collagen fibril cross sections. Fourth row: histograms of collagen fibril diameters*.

**Table 3.**
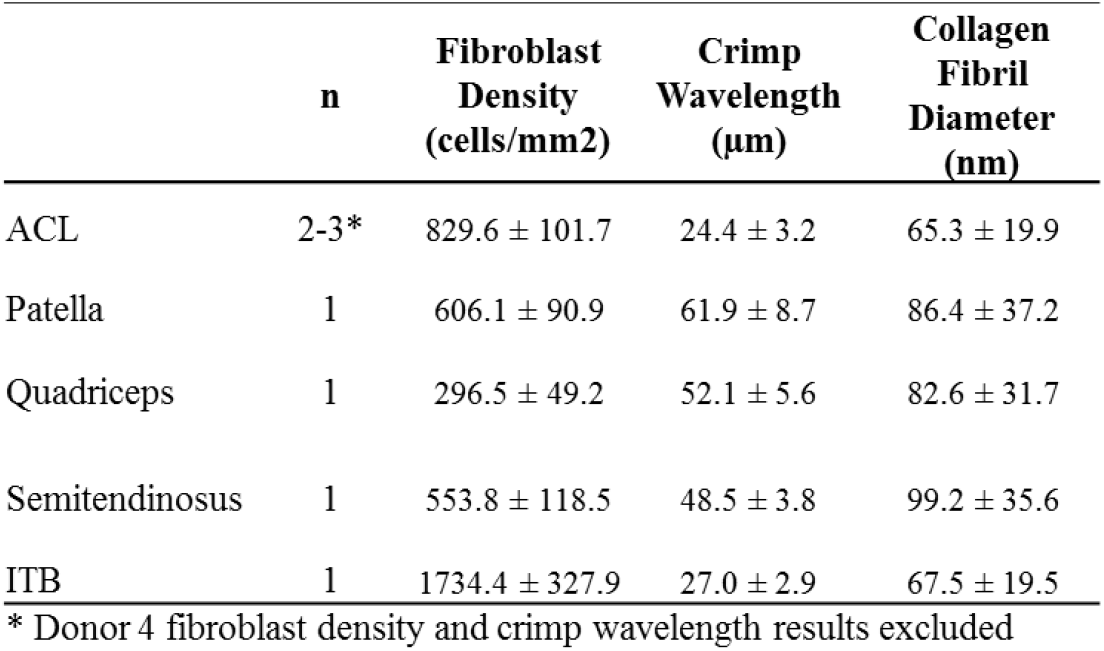
Microstructural analysis results (mean ± standard deviation) for pediatric ACL and graft candidate tendons

| | n | Fibroblast Density (cells/mm<sup>2</sup>) | Crimp Wavelength (μm) | Collagen Fibril Diameter (nm) |
| --- | --- | --- | --- | --- |
| ACL | 2-3* | 829.6 ± 101.7 | 24.4 ± 3.2 | 65.3 ± 19.9 |
| Patella | 1 | 606.1 ± 90.9 | 61.9 ± 8.7 | 86.4 ± 37.2 |
| Quadriceps | 1 | 296.5 ± 49.2 | 52.1 ± 5.6 | 82.6 ± 31.7 |
| Semitendinosus | 1 | 553.8 ± 118.5 | 48.5 ± 3.8 | 99.2 ± 35.6 |
| ITB | 1 | 1734.4 ± 327.9 | 27.0 ± 2.9 | 67.5 ± 19.5 |

\* Donor 4 fibroblast density and crimp wavelength results excluded

Polarized light analysis of crimp morphology revealed interesting similarities and differences between the ACL and the knee tendons of interest (Figure 5). In all of the samples studied, crimp wavelengths appeared extremely uniform, with low amounts of intrafiber variation within the same specimen. The patellar tendon, quadriceps tendon, and semitendinosus tendon displayed longer crimp lengths compared to the ITB and ACL samples (Table 3). Average crimp wavelength for the ACLs from two separate donors was 24.4 ± 3.2 μm.

Results for specimen collagen fiber diameters were similar to that for the crimp wavelengths, with the ITB being most similar to the ACLs. Fibrils in the patellar tendon trended towards smaller diameters, while those in the semitendinosus trended towards larger diameters. The fibrils in the quadriceps tendon were bimodally distributed between large and small diameters. Average fibril diameter for the ACLs from three separate donors exhibited a distinctly unimodal distribution profile centered around a mean of 69.7 ± 20.3 nm (Supplemental Figure 2).

## DISCUSSION

Little is known about the mechanical behavior and microstructural properties of pediatric ACLs and the periarticular tissues most commonly utilized for its reconstruction. Result from this study demonstrated that mechanical properties were considerably weaker than what has been documented for the same structures in healthy adult populations (Figure 6). Interestingly, there are also relative differences in mechanical properties between the native ACL and autograft tendons in adults. For example, all three graft tissues have been shown to exhibit ultimate tensile loads and stiffnesses that are higher than that reported for the native ACL.^18,20,32,34,51,56^ Adult hamstrings tendons seem to possess the strongest mechanical properties and have previously been shown to exhibit significantly higher elastic modulus (1036±312 MPa) and ultimate stress values (120.1±30.0 MPa) than other graft candidates, including the patellar tendon (417±107 MPa, 76.2±25.1 MPa).^48^ This is in agreement with our data for a pediatric population, which showed that the semitendinosus tendons and ITBs are stronger and less compliant than the quadriceps or patellar tendons. We also found relatively large subject-subject variability in the mechanical data for our pediatric specimens, especially for the quadriceps tendon. This could be attributed to donor factors such as age, sex, BMI, and physical activity level, which we could not control for due to the rarity of the donor cohort. Large variations in mechanical properties of adult tendons and ligaments have been previously reported in the literature^48^ and has been further attributed to individual biological variation in collagen fiber maturity, fiber alignment, interfiber cross-linkages and fiber-matrix interaction.^33^

**Figure 6:**
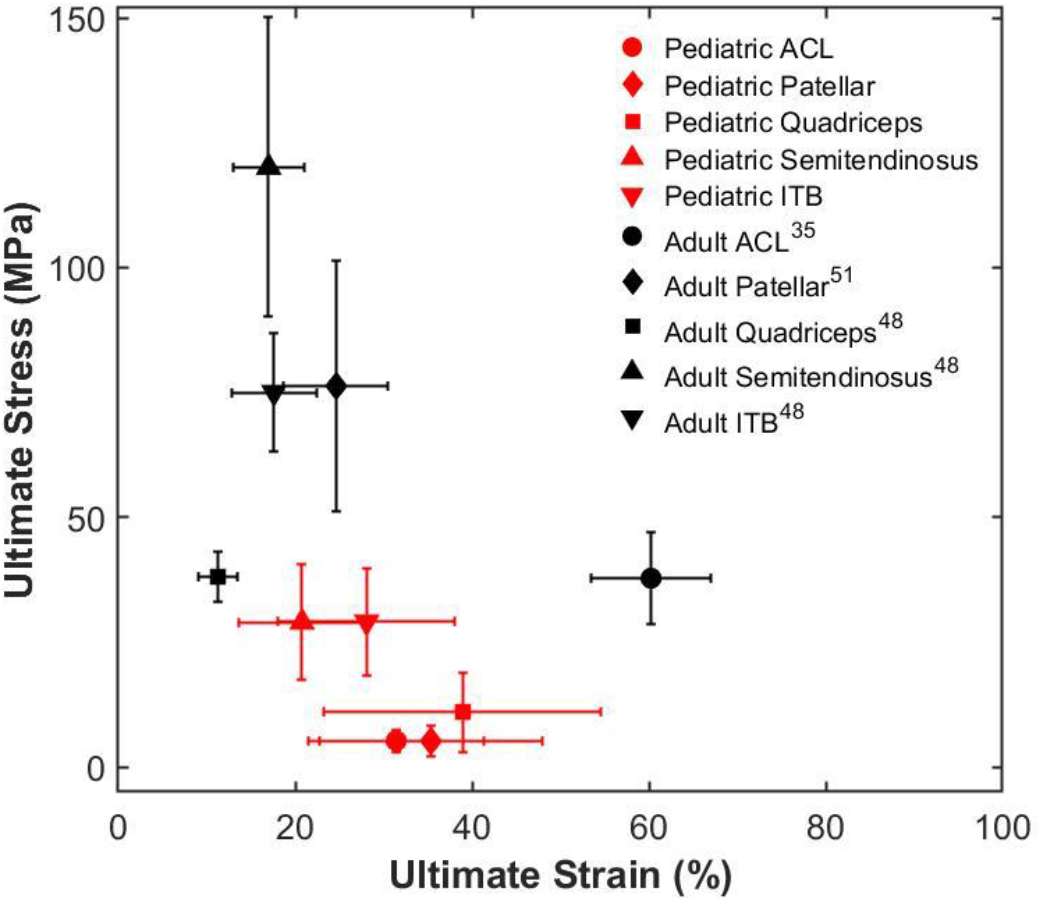
*Ultimate stress-strain plot for the data presented by the current study (non-filled points) compared to data reported in the adult population (filled points). Data points represent the average ultimate stress and ultimate strain and the error bars indicate the standard deviation*.

Functional properties in tendons and ligaments have been shown to be influenced by the morphology of collagen fibrils. Examination of the microstructure in samples harvested from the same knee of one donor revealed distinct differences between the pediatric ACL and the four candidate graft tendons. Crimp morphology has been linked to tendon mechanical function and there is evidence demonstrating that collagen crimp characteristics are strongly tendon-type specific.^50^ Collagen crimps function as a buffer to provide immediate longitudinal elongation in response to load and the non-linear toe region of the stress-strain curve represents the extension of these crimps (Figure 3B).^26^ Past the transition point and into the linear region, the crimps are fully recruited and resistance is provided by the stretching of the collagen triple helix and the cross linkages between these helices. In tendons and ligaments, higher crimp frequency likely provides a protection to the structure during sudden increases in intrinsic forces produced by muscular contractions and compressive joint loads.^12,45^ Results from the current study show that patellar, quadriceps, and semitendinosus tendon samples possessed a lower crimp frequency compared to the ITB and the ACL. This may help to explain the slightly more prolonged toe region for the pediatric ACL (Table 2). The behavior of the toe and transition region is often neglected in the literature, yet it is important to document because ligament strains during activities of daily living and rehabilitation occur within this region.^5,8^ Beynnon & Fleming demonstrated in an *in vivo* study that the strains in the mid substance of the ACL rarely exceed 4% in both weight-bearing and non-weight bearing activities involving varying degrees of knee extension.^5^ Using a mouse model, Miller et al. found that developmentally younger tendons required longer exposure to mechanical load before structurally responding through uncrimping, resulting in higher transition strains.^33^ Interestingly, the pediatric ITB did not exhibit toe region properties similar to the ACL, which demonstrates that the uncrimping of collagen fibers is not the only mechanism driving toe-region properties. For example, collagen fiber crimp frequency has been found to be dependent upon the number of preconditioning cycles that are applied^33^ and it has also been suggested that crimp frequency is heterogeneous along the length of a tendon.^19,52^

The patellar, quadriceps, and semitendinosus tendons showed wider distributions of collagen fibril diameters than the ITB and ACL, which appeared to be predominantly composed of small-diameter fibrils (Figure 5). Mean fibril diameters for the patellar, quadriceps, and semitendinosus tendons were similar to the values reported in the literature for adult samples.^1,17,53^ The mean fibril diameter for the pediatric ACL was considerably smaller than what has been reported for adults.^17^ Little to no information exists in the literature regarding collagen fibril ultrastructure for adult human ITBs. When considering animal models, Qu et al. found significantly higher collagen fibril density and smaller fibril diameters in immature bovine ACLs (102 ± 11.6 nm) compared to a mature group (124.1 ± 22 nm).^42^ They also observed a unimodal distribution of fibril diameters in the immature group and a 3-fold increase in variance in the mature group. It is possible that the ACL is composed of immature networks of collagen fibers for a longer period of time than the periarticular knee tendons.

Collagen fibril diameter size may be reflective of functional adaptations to physiological loads. Fibrils have been shown to be smaller and unimodally distributed at birth, becoming larger and bimodally distributed at maturity.^37,38^ The existence of a true correlation between collagen fibril size and higher stiffness is controversial, but it is a likely contributor. Parry et al. posited that the bimodal distribution of fibril size was optimized so that larger fibrils provided high tensile strength and the smaller fibers provided resistance to creep by increasing the density of interfibrillar electrostatic interactions.^39^

This study has several limitations. Most notably, the sample size in this experiment was low. Due to the rarity of these donors and the use of the specimens across several research labs, this drawback is currently unavoidable. It was difficult to make direct comparisons between the current study and previous studies with adult cohorts. Differences in donor age, sex, physical condition, skeletal maturity, as well as graft size, ramp to failure strain rate, graft construct configuration, method of CSA calculation, and clamping technique can all confound results.^8,56^ The strength of tendons and ligaments has been shown to be dependent on strain rate.^6,24^ To minimize viscoelastic effects, the ramp to failure strain rate was kept at 0.03%/s, which was deliberately chosen to capture toe region properties. Previous studies have utilized strain rates as high as 100%/s to simulate the fast strain rate in accidents and acute injuries.^6,21,35^ This study characterized the mechanical behavior of the tissue at the mid-substance (ACL, patellar tendon, quadriceps tendon) or distal third substance (semitendinosus tendon, ITB) of the tendon or ligament with uniaxial *ex vivo* testing. Many factors that are known to contribute to the strength and durability of an ACLR graft *in vivo*, such as post-operative ligamentization and ligament anchoring,^48^ were not considered. In addition, the anatomical direction of tension is not always axially directed. The ACL is structurally designed to withstand multiaxial stresses that can vary in direction and magnitude along the length of the tissue, which may lead to changes in mechanical and microstructural properties. Many studies have looked at the regional variation in tendon and ligament properties in adults, and this may be a direction for future research in the pediatric population ^2,4^.

### Clinical Implications

Surgical management of ACL deficiency in pediatric patients is complex and many factors can contribute to graft selection for reconstruction. Optimizing ACLR in the pediatric patient population requires a more thorough understanding of anatomical graft placement, the mechanical properties of candidate grafts, the mechanical behavior and strength of anchoring techniques, and the biological processes that occur during graft incorporation. While the mechanical properties of the pediatric patellar tendon were almost identical to that of the pediatric ACL, the semitendinosus and ITB were considerably stronger and could resist loads that make the ACL prone to reinjury or caused the injury in the first place. Another important consideration is that the mechanical properties of ACLR grafts have been demonstrated to decrease during the remodeling process and never return to normal,^43^ making these even more attractive graft candidates.

Double-bundle reconstruction has been used in an effort to improve knee stability and restore the biomechanical characteristics of the intact ACL. Some previous work has identified higher risk of physeal injury with double bundle techniques in those with open physis, and thus, these procedures may not be ideal in younger patients with significant growth remaining.^46^ Younger children often possess small hamstrings tendons that have insufficient graft diameter, yield smaller final graft sizes, and ultimately increase the likelihood of revision. In cases whereby smaller hamstring graft sizes are noted intraoperatively there is a recent trend toward increasing the number of strands used via a folding technique which increases graft diameter.^31^ This study assessed the mechanical properties of the distal third substance of non-folded semitendinosus tendon and does not consider the mechanical implications of the folding techniques that are used for multi-bundle reconstructions (such as 4, 6, 8 strand constructs), however this would be an excellent direction for future research.

The incidence of pediatric ACL injuries has increased significantly over the past several decades as youth participation and specialization in sports has increased. Despite this, there is a dearth of information on the mechanical and microstructural properties of pediatric knee tendons and ligaments in the literature. Further optimization of the pediatric ACLR procedure such that the surrogate graft is the most appropriate mechanical and biologic option for the patient will ultimately improve surgical outcomes. This study utilized a small and extremely rare sample of pediatric cadaveric knees to develop a comprehensive suite of data on the pediatric ACL and the most common tendons used for reconstruction in the pediatric knee. Our data showed that different autografts possess distinct mechanical properties that often differ markedly from the ACL. This data can be used to inform the selection of grafts for reconstruction, the design and fabrication of synthetic constructs, and also to develop constitutive musculoskeletal models that can be applied to clinically relevant loading conditions that are difficult to test with bench-top experiments alone.

## ACKNOWLEDGMENTS

The authors thank Allosource for the donation of the specimens used in this study. This study was supported by the Penn Center for Musculoskeletal Disorders Histology Core (NIH P30-AR06919) and by the Electron Microscopy Resource Laboratory of the University of Pennsylvania.

